# A genome-scale metabolic network model and machine learning predict amino acid concentrations in Chinese Hamster Ovary cell cultures

**DOI:** 10.1101/2020.09.02.279687

**Authors:** Song-Min Schinn, Carly Morrison, Wei Wei, Lin Zhang, Nathan E. Lewis

## Abstract

The control of nutrient availability is critical to large-scale manufacturing of biotherapeutics. However, the quantification of proteinogenic amino acids is time-consuming and thus is difficult to implement for real-time *in situ* bioprocess control. Genome-scale metabolic models describe the metabolic conversion from media nutrients to proliferation and recombinant protein production, and therefore are a promising platform for *in silico* monitoring and prediction of amino acid concentrations. This potential has not been realized due to unresolved challenges: (1) the models assume an optimal and highly efficient metabolism, and therefore tend to underestimate amino acid consumption, and (2) the models assume a steady state, and therefore have a short forecast range. We address these challenges by integrating machine learning with the metabolic models. Through this we demonstrate accurate and time-course dependent prediction of individual amino acid concentration in culture medium throughout the production process. Thus, these models can be deployed to control nutrient feeding to avoid premature nutrient depletion or provide early predictions of failed bioreactor runs.

## Short Communication

Chinese Hamster Ovary (CHO) cells are widely used to manufacture complex biotherapeutic molecules at large scales. Industrial bioprocesses ensure high product yield and quality by maintaining favorable growth conditions in cell culture environments, which requires careful monitoring and control of nutrient availability. Chemically-defined serum-free media can contain dozens or >100 components (Ritacco et al., 2018), but key nutrients include proteinogenic amino acids, which are direct substrates and regulators (Duarte et al., 2014; Fomina◻Yadlin et al., 2014) of proliferation and protein synthesis. Unfortunately, conventional methods for amino acid quantification based on liquid chromatography and mass spectrometry are time-consuming and difficult to use for decision making and control of cell culture. Alternate spectroscopic approaches have been sensitive to a limited number of amino acid species (Bhatia et al., 2018). Here we present a computational method to forecast time-course amino acid concentrations from routine bioprocess measurements, facilitating a timely and anticipatory control of the bioprocess (Fig. 1).

**Figure 1:**
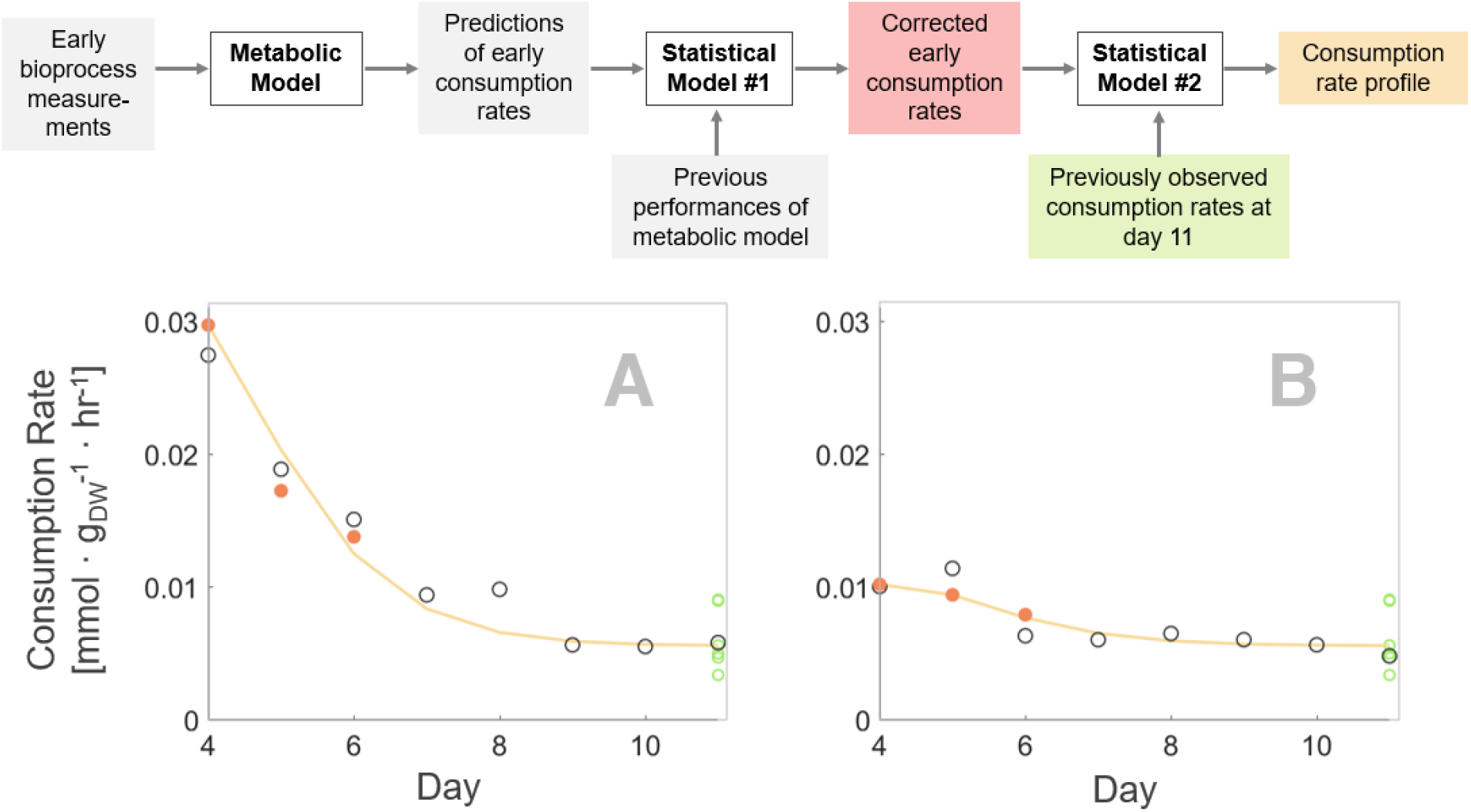
Overview of method. A novel combination of a metabolic and statistical models forecast the time-course amino acid consumption profiles in CHO cell cultures. A workflow of the prediction procedure is provided; key data are color-coded and visualized in plots A & B by the same color. First, a metabolic model predicts amino acid consumption rates for days 4-6 based on routine bioprocess measurements such as viable cell density and glucose uptake rate. Then, a statistical model refines these predictions (red) by considering the metabolic model’s previous performances. Based on these predictions from early culture days, a second statistical model predicts the complete consumption profile (orange). The model references asymptotic behavior of previous consumption profiles as priors (green). The predicted consumption profiles agreed well with experimental data (black empty markers). The two plots show distinct leucine consumption profiles from CHO clones C2 and Z3 (Fig. S1) with disparate early consumption patterns. A more detailed workflow can be found in Fig. S6.

At the foundation of our method is a genome-scale metabolic network model, which accounts for the complex conversion from media nutrients to biomass and recombinant protein production. Such models have been increasingly utilized for CHO cells (Hefzi et al., 2016; Calmels et al., 2019; Huang & Yoon, 2020) and bioprocess applications (Sommeregger et al., 2017), such as predicting clonal performances (Popp et al., 2016), identifying metabolic bottlenecks (Zhuangrong & Seongkyu, 2020), and optimizing media formulation (Fouladiha et al., 2020; Traustason et al., 2019). Metabolic network models can also estimate amino acid uptake rates necessary to experimentally support observed proliferation and productivity (Chen et al., 2019). However, several challenges have limited their practical application.

First, metabolic network models are typically highly complex but under-constrained, and therefore are easy to overfit. This is mitigated by training the model on a variety of bioprocess conditions and metabolic phenotypes. Second, metabolic network models assume that cells operate at some metabolic optimum, and thus tend to describe an idealized metabolism specifically fit to the assumed objective, e.g., biomass production (Feist & Palsson, 2010; Szeliova et al., 2020), minimization of redox (Savinell & Palsson, 1992). Third, for the present purpose, these models need to predict amino acid consumption fluxes, typically on the order of 10^−3^ mmol·g_DW_^−1^·hr^−1^(see Methods), from input data that are multiple magnitudes larger, such as growth rate and glucose consumption (10^−1^ to 10^−2^ mmol·g_DW_^−1^·hr^−1^). The preceding two challenges increase prediction error. Lastly, metabolic network models assume a steady state, which reduces the range of forecast. Typically, input data from one day are used to make predictions for the same day. However, such predictions cannot be extended to multiple days or subsequent culture phases, as cross-temporal shifts in metabolism would violate the steady state assumption. In summary, model predictions of amino acid concentrations can be overfit, ideal, and near-sighted – all of which dilutes their practicality for industrial bioprocess control. Here we demonstrate that these weaknesses can be addressed in a data-driven manner by coupling a metabolic network model with machine learning.

We developed this hybrid approach on a diverse set of 10 CHO clones with different growth and productivity profiles from two different fed-batch production processes. These CHO clones were subject to different bioprocess conditions and recombinant antibody identities (see Methods), resulting in a variety of phenotypes and productivity performances (Fig. S1). For example, several high-performing clones were exceptionally proliferative or productive, suggesting an efficient conversion from nutrients to biomass or recombinant protein product. Other clones performed these conversions at lower rates, suggesting attenuated metabolic activity or inefficient resource utilization. The CHO cells adjusted their nutrient uptake according to these various metabolic phenotypes, leading to diverse amino acid consumption patterns (Fig, S2). For example, the consumption of glucose and serine differed by several fold across conditions and time. Furthermore, different clones varied in their consumption or secretion of key metabolites such as lactate, alanine, glycine, and glutamine.

We sought to predict these diverse consumption behaviors using a tailored model of CHO metabolism (Table S1, S2). As input information, we utilized the following routinely measured industrial bioprocess data: (1) viable cell density and titer measurements, from which growth rate and specific productivity are calculated (Methods, equation 1), and (2) bioreactor concentrations of glucose, lactate, glutamate and glutamine, from which their respective consumption rates are calculated. These measurements were used as boundary conditions by constraining the fluxes of biomass production, recombinant protein synthesis and consumption of the four metabolites to observed values. Subsequently, we used Markov chain Monte Carlo sampling of metabolic fluxes (Schellenberger et al., 2011) to sample the range and magnitude of all reaction fluxes to calculate the likely uptake fluxes of the remaining 18 proteinogenic amino acids (see Methods). These predictions were applied to the CHO clones across 8 days of a 12-day production run (days 4 to 11), resulting in a total of 80 individual predictions.

Predictions from the metabolic model agreed well with experimental measurements. Prediction errors were small compard to the scale of input data (Fig. 2A), suggesting that metabolic models can describe the conversion from nutrients to biomass and recombinent proteins. However, the model also underestimated consumption rates for almost all amino acids (Fig. 2B, *x*-axis), on average by about half a fold. This is likely because the model doesn’t consider certain metabolic inefficiencies – e.g. futile cycles or cytotoxic byproduct synthesis (Mulukutla et al., 2017).

**Figure 2:**
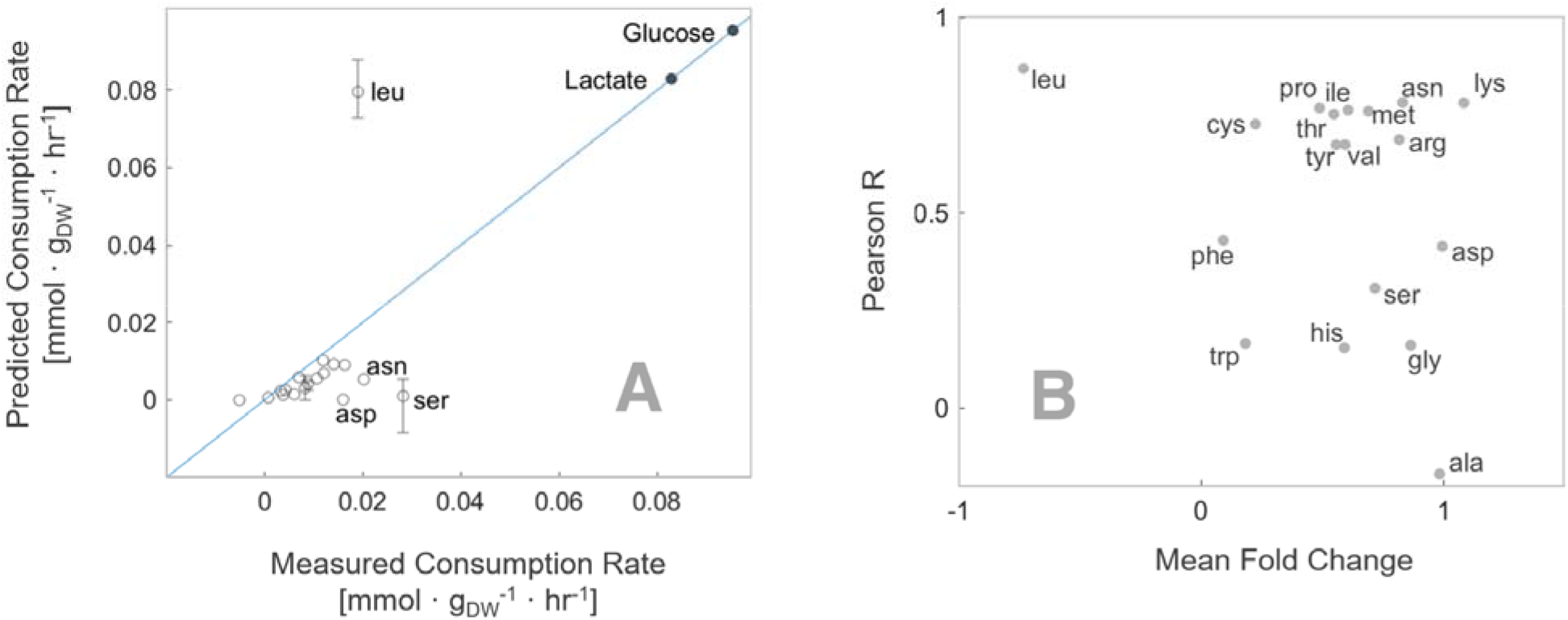
Metabolic network model estimates amino acid consumption rates. (a) Model predictions compared well to experimental observations, given the scale of input data such as the consumption rates of glucose and lactate (upper right, filled circles). (b) However, the fold change between model predictions and experimental measurements could be significant for several amino acids (*x*-axis). Fortunately, the relatively high linear correlation between predictions and measurements (*y*-axis) suggests that predictions could be improved empirically.

Notably, the predicted consumption rates correlated well with measurements for many amino acids (Fig. 2B, *y*-axis). Therefore, we constructed a series of linear regression models to ‘correct’ the metabolic model predictions, using the predicted values and growth rate as explanatory variables (Methods, equation 2). This substantially improved predictions for most amino acids (Fig. S4). As exceptions, predictions for alanine and glycine did not sufficiently improve due to their high fold change error and low correlation to experimental measurements (Fig. 2B). These amino acids are non-essential and can be synthesized from glucose cost-efficiently. Therefore, their consumption may be regulated distinctly and more independent from growth requirements, as observed previously in other organisms (M. Zampieri et al., 2019). Indeed, alanine and glycine were the only two amino acid species that were variously consumed and secreted in significant amounts (Fig. S2). In short, the investigated CHO cells seem to consume them in a ‘less ideal’ manner than other amino acids.

Overall, our hybrid modeling approach estimated most amino acid consumptions well at a small timescale (1 day), when the steady state assumption holds true. This assumption is not valid at larger timescales of multiple days, where nutrient consumption declines asymptotically as cellular metabolism shifts from exponential growth phase to stationary phase. We addressed this limitation by modeling the multi-phase consumption profile with an exponential decay function (Methods, equation 3; Figure S5). Specifically, we first predicted amino acid consumption rates of several early culture days as heretofore described (Fig. 1, red datapoints). Then, these datapoints were used to fit an exponential decay function that describes the entire consumption profile, including later culture days (Fig. 1, orange line).

Our approach accurately predicted daily consumption rates for each amino acid excluding alanine and glycine (Fig. 3A). This included amino acids that are highly abundant in recombinant antibodies (e.g. serine, valine, and leucine) (Fan et al., 2015), or that complicate media formulation due to low solubility (e.g. tyrosine). We also estimated the total amounts of amino acid consumed over the 8 culture days to within 86% of experimental values (Fig. 3B). These results highlight the method’s value in monitoring and forecasting the bioreactor environment.

**Figure 3:**
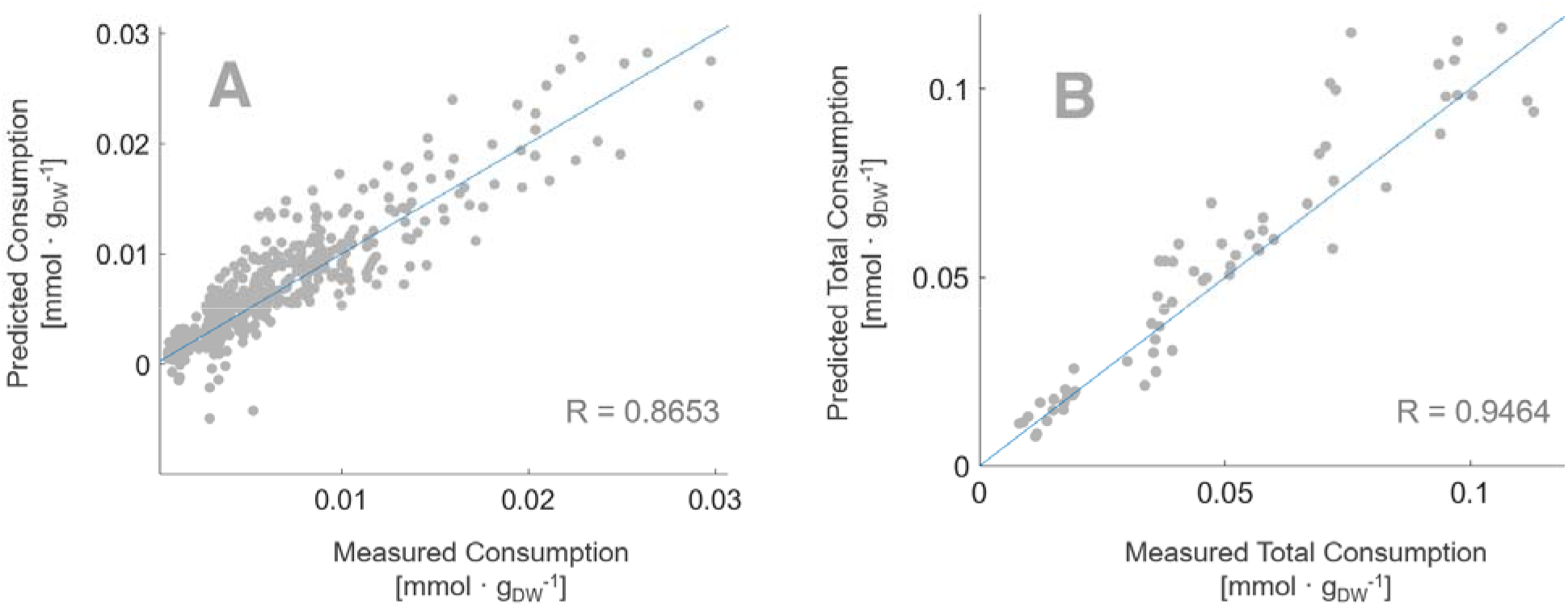
Statistical model forecasts consumption profiles. (A) For validation, the daily consumption rates were calculated for 16 amino acids (excluding alanine and glycine). On average, the predicted values agreed with experimental measurements to within 83%. (B) Then, the total amount of amino acid consumed across the investigated culture period were calculated by summing the daily consumption rates, which agreed with experimental measurements to within 86% on average. This suggests that the method can track and forecast the bioreactor environment.

In summary, the presented modeling workflow forecasted the entire amino acid consumption profile from early bioprocess measurements, facilitating anticipatory and *in situ* control of bioreactor nutrient availability. This was realized by a novel combination of metabolic and statistical models. A metabolic network model estimated amino acid uptake rates necessary for observed proliferation and productivity, assuming an ideally efficient metabolism and steady state conditions. Two subsequent statistical models refined these predictions by offsetting prediction errors empirically and by describing the time-course relationship of individual predictions. These statistical models can easily be adjusted and re-trained for changes in cell-lines or bioprocesses. Our efforts are part of a growing trend of synergizing metabolic network models with machine learning methods (G. Zampieri et al., 2019), and demonstrates the power of hybrid modeling for on-line control of bioprocesses.

## Methods

### Cell culture experiments

Two production fed batch processes were used, Fed batch 1 and Fed batch 2. Both fed batch processes used chemically defined media and feeds over the 12-day cell culture. Fed batch 1 used a glucose restricted fed batch process called HiPDOG (Gagnon et al., 2011). Glucose concentration is kept low during the initial phase of the process, Day 2-7, through intermittent addition of feed medium containing glucose at the high end of pH dead-band and then glucose was maintained above 1.5 g/L thereafter, restricting lactate production without compromising the proliferative capability of cells. In Fed batch 2 a conventional cell culture process was used where glucose was maintained above 1.5 g/L throughout the process.

For both process conditions, bioreactor vessels were inoculated at 2 x 10^6^ viable cells/mL. The following bioprocess characteristics were quantified daily using a NOVA Flex BioProfile Analyzer (Nova Biomedical, Waltham, MA): viable cell density, average live cell diameter and concentrations of glucose, lactate, glutamate, and glutamine. Viable cell density data were converted to growth rates by following equation to be compared to model-predicted growth rates.

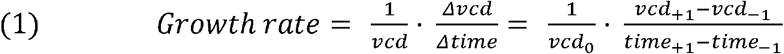

Flash-frozen cell pellets (10^6^ cells) and supernatant (1 mL) were collected from bioreactor runs for each sampling day. Collected samples were sent to Metabolon (Metabolon Inc, Morrisville, NC) for metabolomics analyses. Metabolomics measurements were used as input data to the model by converting their units to model units of mmol per gram of dry weight of cell per hour.

### Metabolic network modeling

We used a previously described metabolic network model that is tailored to the investigated CHO clones (Schinn et al., 2020). Experimental measurements for clone and culture day were used to constrain model reactions for biomass production, monoclonal antibody secretion and consumption of glucose, lactate, glutamate, and glutamine. Then, we computed distributions of likely amino acid consumption rates by stochastically sampling 5000 points within the model’s solution space via a Markov chain Monte Carlo sampling algorithm, as described previously (Nam et al., 2012), using *optGpSampler* (Megchelenbrink et al., 2014) and COBRApy (Ebrahim et al., 2013).

### Statistical methods

For each amino acid, the mean of the sample distribution was interpreted as likely consumption rates predicted by the metabolic model. These predictions were refined and extended by statistical models, as explained below. The modeling workflow is visualized in a detailed diagram (Fig. S6) and demonstrated by sample code (Supplementary Data). Specifically, the statistical models were trained and validated by randomly dividing the 80 observations into two sets, consisting of 48 and 32 observations, respectively. Quantified snapshots of the validation data throughout the analysis workflow are detailed in supplementary tables, from experimental measurement to final model prediction (Table S3, S4, S6, S10); priors and inferences derived from the training data set are also provided (Table S5, S7, S8, S9).

The first statistical model refined metabolic model predictions (equation 2). Growth rate was included as an explanatory variable as it also correlated well with consumption rates of many amino acid species (Fig. S3).

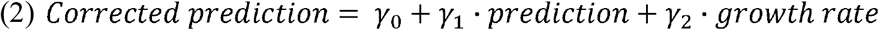

The second statistical model described time-course amino acid consumption by an exponential decay function (equation 3). The coefficient β_0_ represents the minimum consumption rate which the cells asymptotically approach during later stationary phase. First, regression coefficients were calculated from the training dataset to be used as priors and constraints for nonlinear optimization. Specifically, the mean, minimum and maximum values of these training coefficients were used as initial guess values, lower bounds, and upper bounds, respectively.

Then, regression coefficients were fitted to minimize two values: (1) the difference between outputs of equation 2 and equation 3 for early culture days, (2) the difference between fitted β_0_ and previously observed asymptotic values.

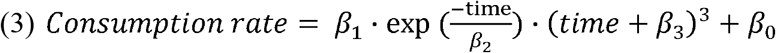

These analyses were carried out and visualized using COBRA Toolbox 2.0 (Schellenberger et al., 2011) in MATLAB R2018b (MathWorks; Natick, Massachusetts, USA)

