## Supplementary Figures for "A genome-scale metabolic network model and machine learning predict amino acid concentrations in Chinese Hamster Ovary cell cultures"

### Slide 1
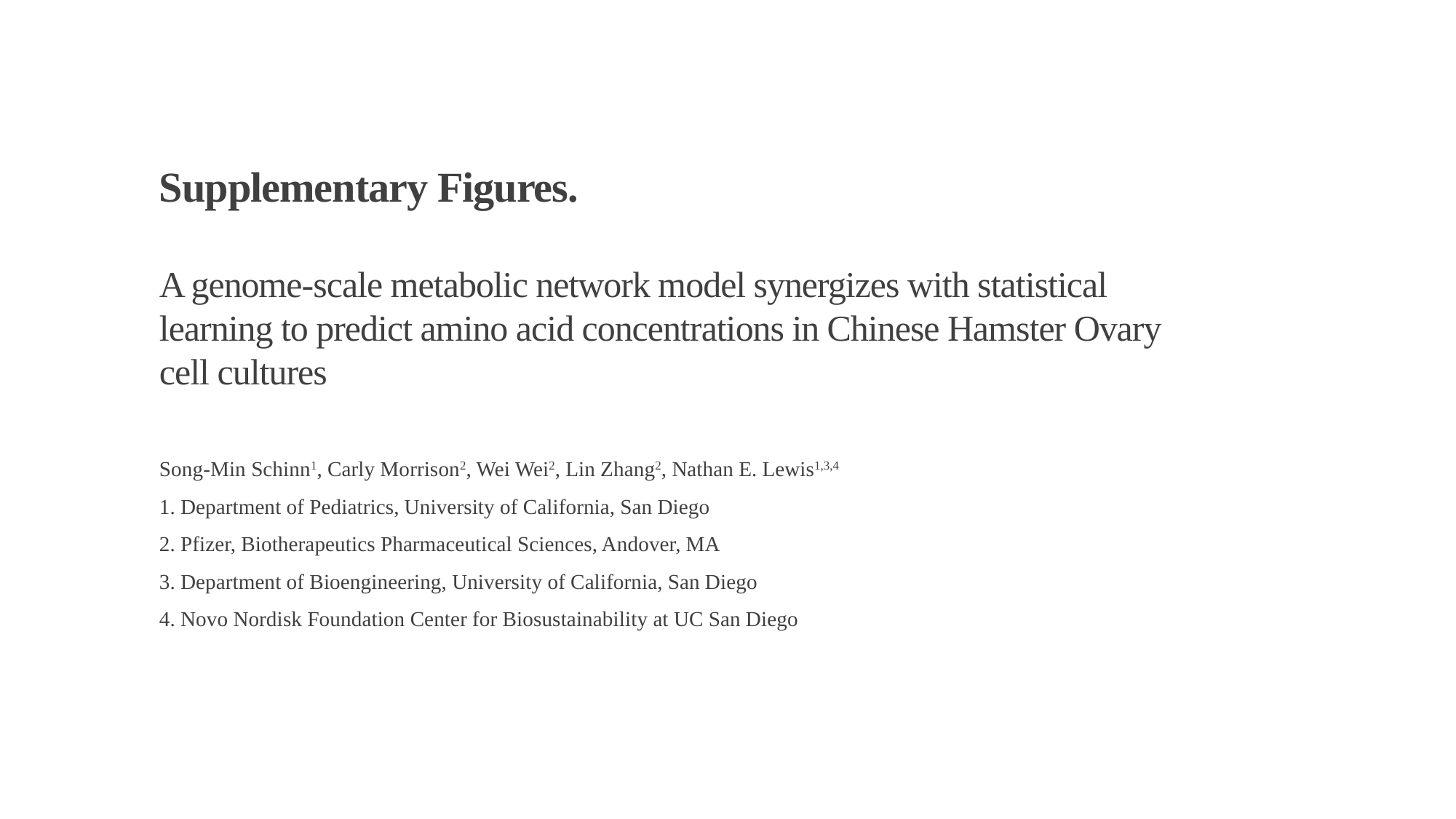

Supplementary Figures.
A genome-scale metabolic network model synergizes with statistical learning to predict amino acid concentrations in Chinese Hamster Ovary cell cultures
Song-Min Schinn^1^, Carly Morrison^2^, Wei Wei^2^, Lin Zhang^2^, Nathan E. Lewis^1,3,4^
1. Department of Pediatrics, University of California, San Diego
2. Pfizer, Biotherapeutics Pharmaceutical Sciences, Andover, MA
3. Department of Bioengineering, University of California, San Diego
4. Novo Nordisk Foundation Center for Biosustainability at UC San Diego

### Slide 2
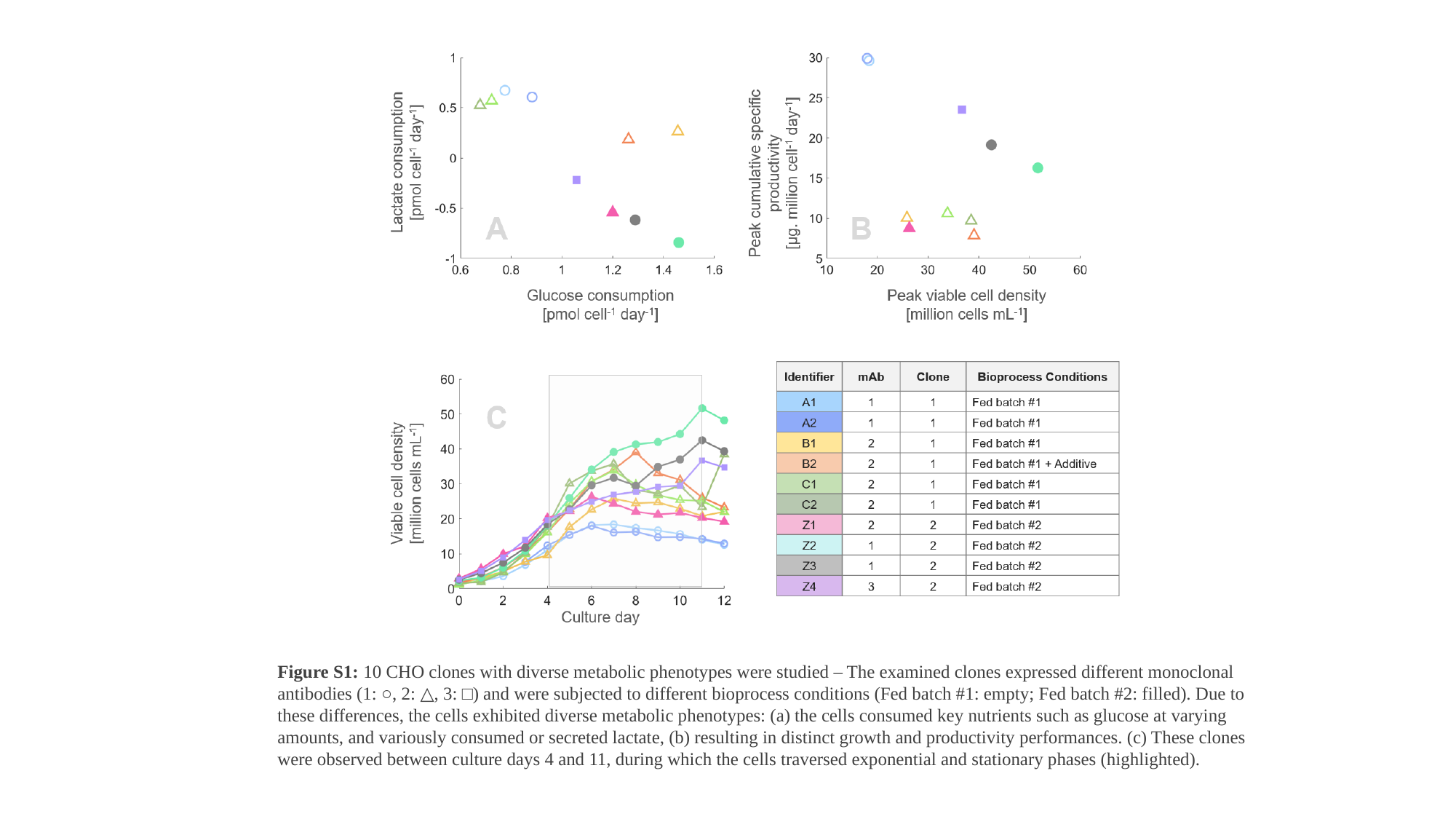

Figure S1: 10 CHO clones with diverse metabolic phenotypes were studied – The examined clones expressed different monoclonal antibodies (1: ○, 2: △, 3: □) and were subjected to different bioprocess conditions (Fed batch #1: empty; Fed batch #2: filled). Due to these differences, the cells exhibited diverse metabolic phenotypes: (a) the cells consumed key nutrients such as glucose at varying amounts, and variously consumed or secreted lactate, (b) resulting in distinct growth and productivity performances. (c) These clones were observed between culture days 4 and 11, during which the cells traversed exponential and stationary phases (highlighted).

### Slide 3
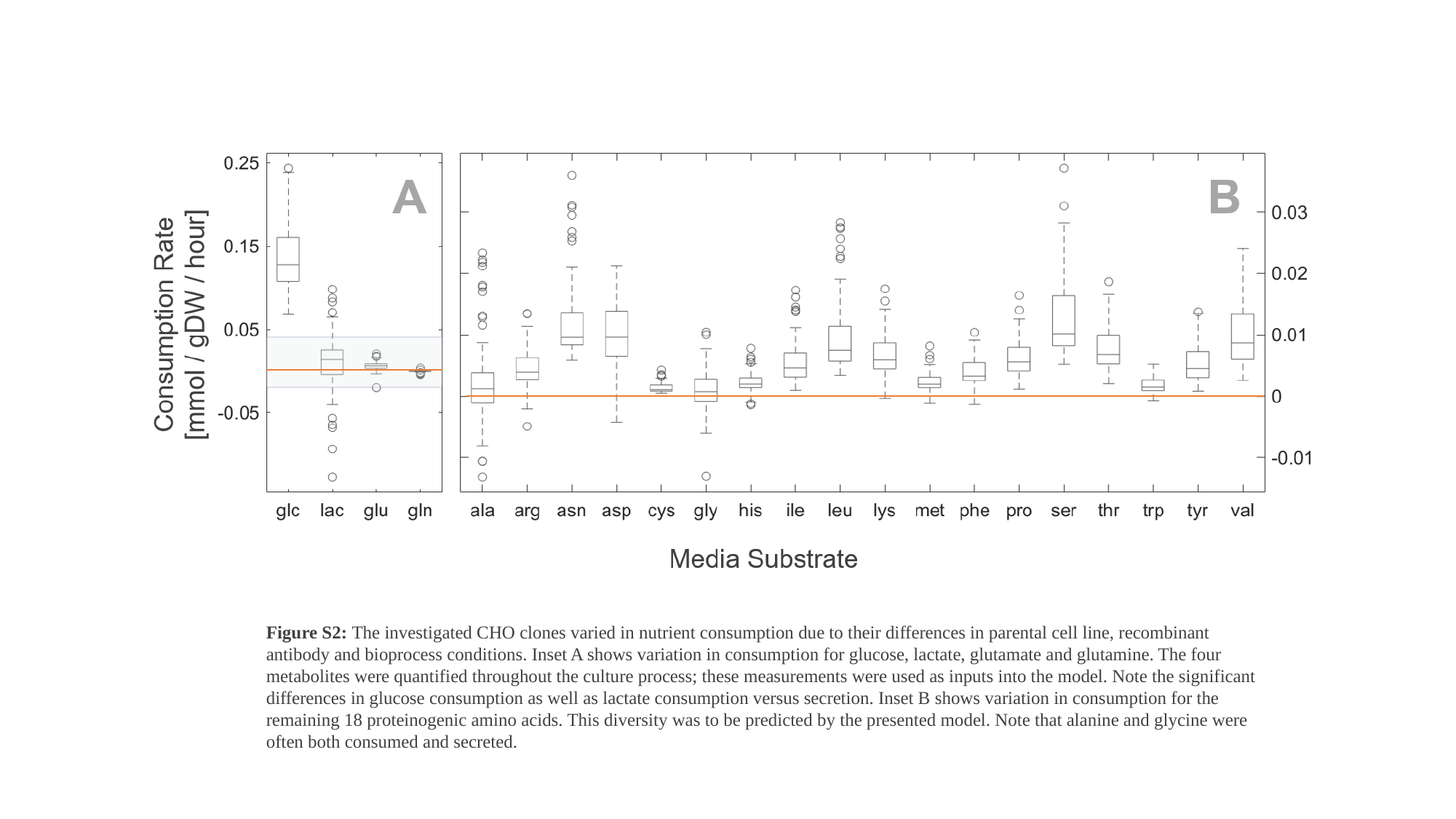

Figure S2: The investigated CHO clones varied in nutrient consumption due to their differences in parental cell line, recombinant antibody and bioprocess conditions. Inset A shows variation in consumption for glucose, lactate, glutamate and glutamine. The four metabolites were quantified throughout the culture process; these measurements were used as inputs into the model. Note the significant differences in glucose consumption as well as lactate consumption versus secretion. Inset B shows variation in consumption for the remaining 18 proteinogenic amino acids. This diversity was to be predicted by the presented model. Note that alanine and glycine were often both consumed and secreted.

### Slide 4
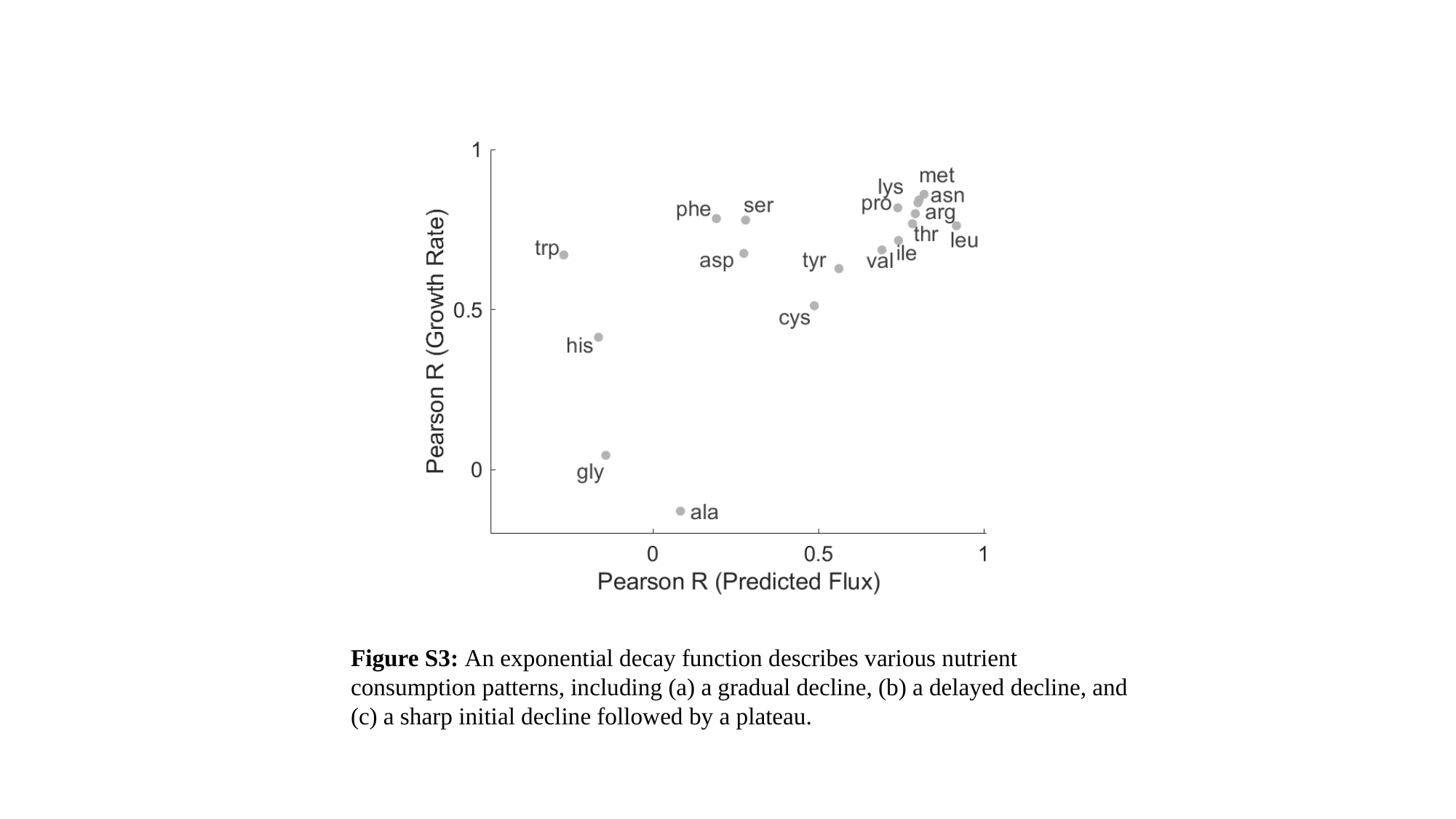

Figure S3: An exponential decay function describes various nutrient consumption patterns, including (a) a gradual decline, (b) a delayed decline, and (c) a sharp initial decline followed by a plateau.

### Slide 5
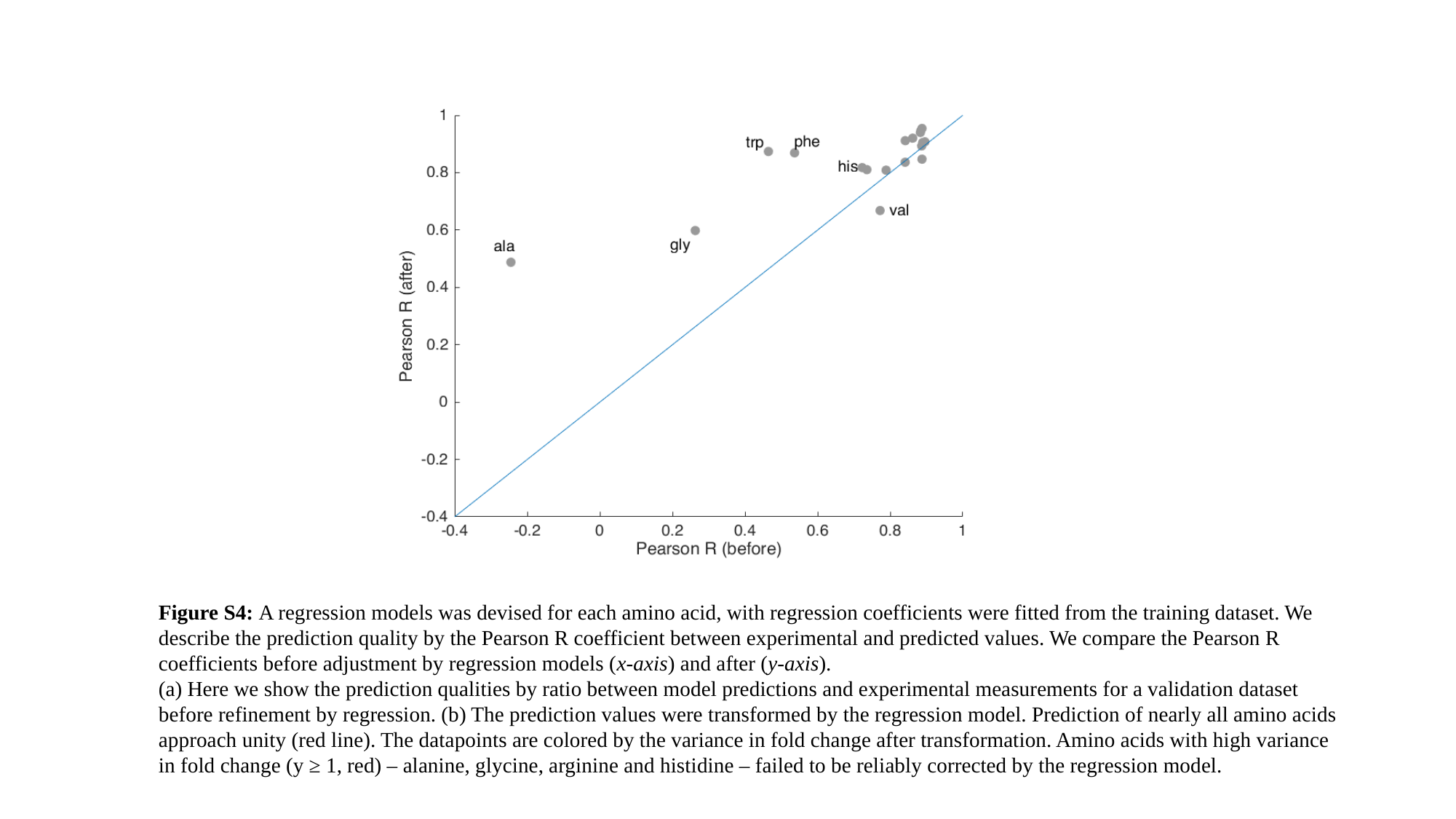

Figure S4: A regression models was devised for each amino acid, with regression coefficients were fitted from the training dataset. We describe the prediction quality by the Pearson R coefficient between experimental and predicted values. We compare the Pearson R coefficients before adjustment by regression models (x-axis) and after (y-axis).
(a) Here we show the prediction qualities by ratio between model predictions and experimental measurements for a validation dataset before refinement by regression. (b) The prediction values were transformed by the regression model. Prediction of nearly all amino acids approach unity (red line). The datapoints are colored by the variance in fold change after transformation. Amino acids with high variance in fold change (y ≥ 1, red) – alanine, glycine, arginine and histidine – failed to be reliably corrected by the regression model.

### Slide 6
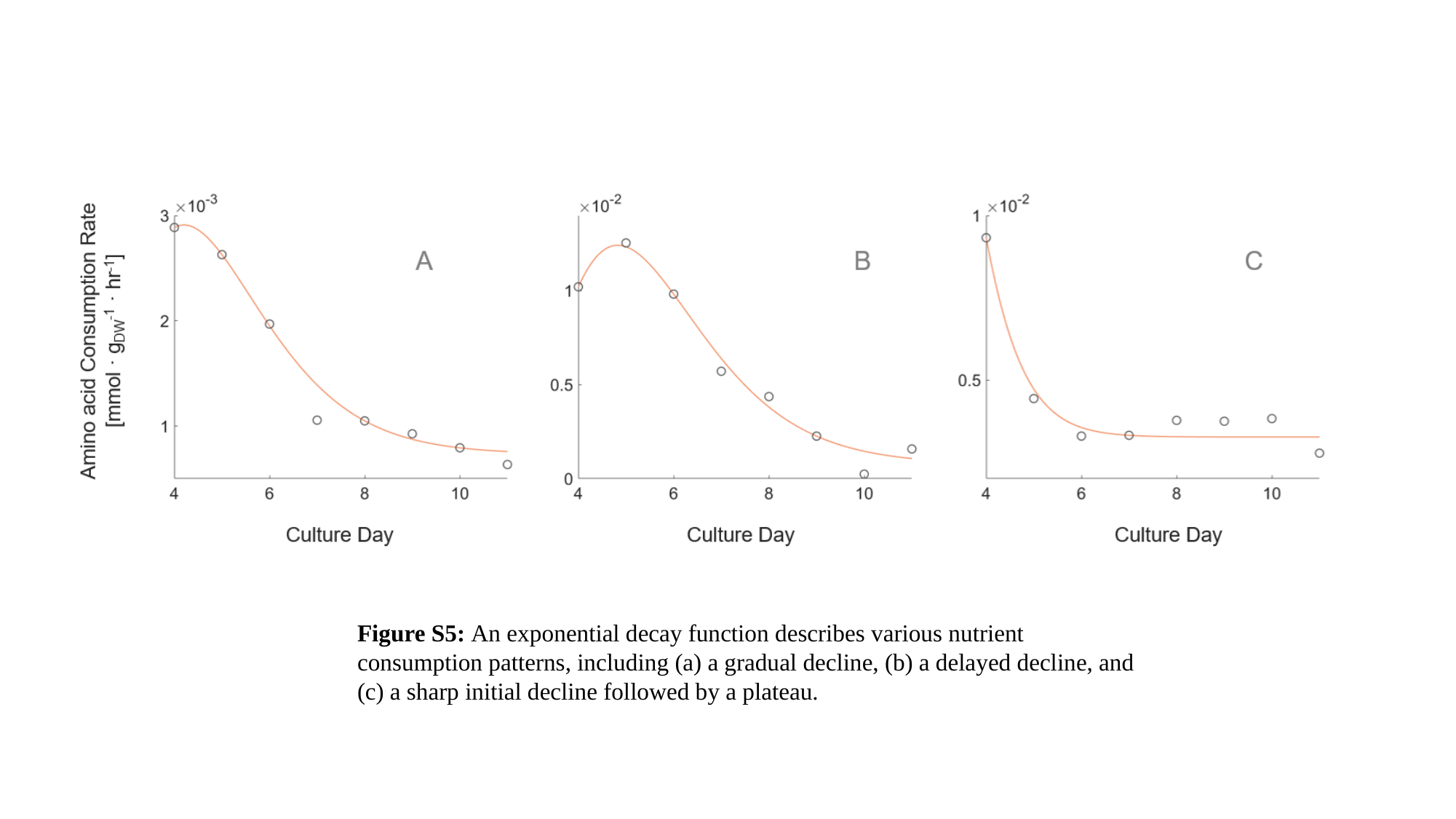

Figure S5: An exponential decay function describes various nutrient consumption patterns, including (a) a gradual decline, (b) a delayed decline, and (c) a sharp initial decline followed by a plateau.

### Slide 7
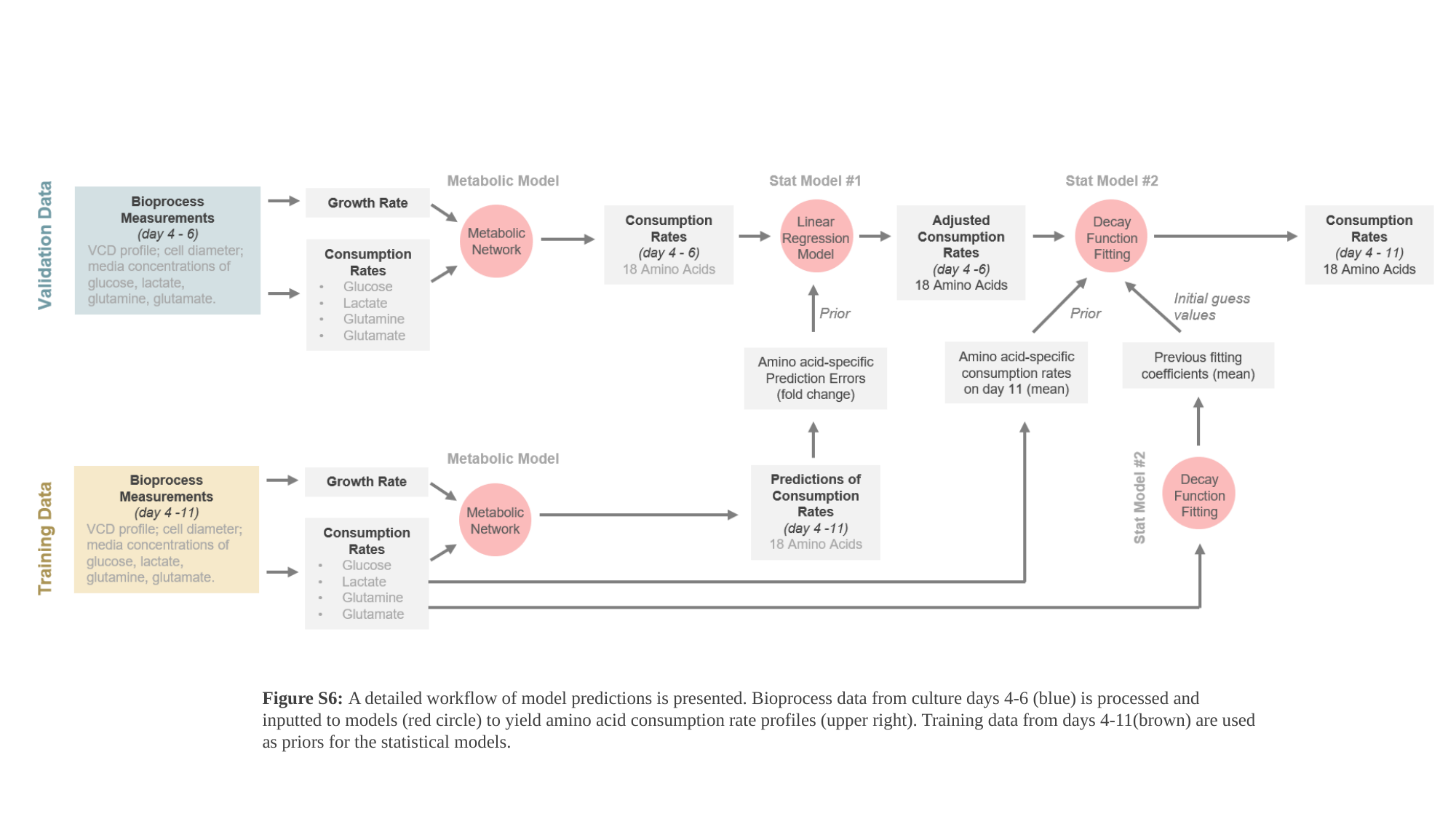

Figure S6: A detailed workflow of model predictions is presented. Bioprocess data from culture days 4-6 (blue) is processed and inputted to models (red circle) to yield amino acid consumption rate profiles (upper right). Training data from days 4-11(brown) are used as priors for the statistical models.
