## Supplementary material for "A genome-scale metabolic network model and machine learning predict amino acid concentrations in Chinese Hamster Ovary cell cultures": Code & Data: Predict_Aminoacid_Profiles.m

```
addpath('./data');
amino_acids ={'leu','tyr','asp','cys','asn','ser','gly','his','thr','arg','ala',...
    'val','met','trp','phe','ile','lys','pro'};
amino_acids = sort(amino_acids);
expm_data = readtable('expm_data.csv');
output1 = readtable('output1.csv'); %this is model output from model #1 (metabolic model)

%% Statistical Model #1
model_2 = readtable('statmodel1_coeffs.csv');
F = [];
for i = 1:length(amino_acids)
    x = output1{:,amino_acids{i}};
    mu = expm_data{:, 'mu'};
    X = [ones(length(x),1) mu x];
    Y = X*model_2{:,amino_acids{i}};
    F = [F, Y];
end
output2 = array2table(F);
output2.Properties.VariableNames = amino_acids;
output2.replicate = expm_data.replicate;
output2.day = expm_data.day;
% These are consumption predictions that are 'corrected' by statistical
% model #1. You can compare this with 'output2.csv' provided in data folder

%% Statistical Model #2
replicates = unique(expm_data.replicate);
trained_coeffs = readtable('trained_coefficients.csv');
consump_day11 = readtable('asymptotes.csv');
earlyday_threshold = 6;
M = [];
for i = 1:length(amino_acids)
    F = [];
    
    for j = 1:length(replicates)
        ix = find(output2.replicate == replicates(j));
        x = output2{ix, 'day'};  
        y = -output2{ix, amino_acids{i}};
        early_y = y(x<=earlyday_threshold); 
        early_x = x(x<=earlyday_threshold);
        
        % fit with priors from Training Data (simple)
        ix = find(strcmp(trained_coeffs.met, amino_acids{i}));
%         ix = intersect(ix,  find(trained_coeffs.repl == valid_repls(j)));
        my_coeffs = trained_coeffs{ix,contains(trained_coeffs.Properties.VariableNames, 'coeff')};
        lb = min(my_coeffs); 
        ub = max(my_coeffs);
        init_guess_vals = mean(my_coeffs,1);
        late_flux = mean(consump_day11{:, amino_acids{i}});
        [newx, newy, coeffs3] = fit_w_constraints(early_x, early_y,late_flux,init_guess_vals, lb, ub);
        F = [F; newy];
    end
    M = [M, F];
end
output3 = array2table(M);
output3.Properties.VariableNames = amino_acids;
output3.replicate = expm_data.replicate;
output3.day = expm_data.day;
% These are the consumption profiles predicted by statistical model #2. You
% can compare this with 'output3.csv' or Supplementary Table 10.

%% Visualize
figure()
amino_acid_of_interest = 'leu';
for j = 1:length(replicates)
    ix = find(expm_data.replicate == replicates(j));
    time = expm_data{ix, 'day'};
    measured = -expm_data{ix, amino_acid_of_interest}; 
    predicted = -output2{ix, amino_acid_of_interest}; 
    profile = output3{ix, amino_acid_of_interest};
    
    subplot(2,2,j); hold on;
    plot(time, measured, 'o')
    plot(time([1:3]), predicted([1:3]), '^')
    plot(time, profile, '-')
    xlabel('day')
    ylabel('consumption rate') % unit:  [mmol/gdw/hr]
end

%% --- local function

function [newx, newy, coeffs] = fit_w_constraints(x, y, y11, init_guess, lb, ub)
% regression via 'fmincon'
options = optimoptions('fmincon');
options.MaxFunctionEvaluations = 9000;
options.MaxIterations = 6000;

%       y = c*exp(-x/b)*(x+a)^3+d
n = 3;
errorf = @(C) ...
    sum(abs( (    C(3).*exp(-x./C(2)).*(x+C(1)).^n + C(4) - y     ))) + ...
    sum(abs( (    C(3).*exp(-11./C(2)).*(11+C(1)).^n + C(4) - y11     ))) ;

C = fmincon(errorf, init_guess, [], [], [], [], lb, ub, [] , options);
a = C(1); b = C(2);   c = C(3); d = C(4);
newx = [4:11]';
newy =  c.*exp(-newx./b).*(newx+a).^n+d;
coeffs = [a b c d];

end
```
